# Displacement statistics of unhindered single molecules show no enhanced diffusion in enzymatic reactions

**DOI:** 10.1101/2021.07.20.452795

**Authors:** Alexander A. Choi, Ha H. Park, Kun Chen, Rui Yan, Wan Li, Ke Xu

## Abstract

Recent studies have sparked heated debate over whether catalytical reactions would enhance the diffusion coefficients *D* of enzymes. Through high statistics of the transient (600 μs) displacements of unhindered single molecules freely diffusing in common buffers, we here quantify *D* for four highly contested enzymes under catalytic turnovers. We thus formulate how precisions of better than ±1% may be achieved for *D* at the 95% confidence level, and show no changes in diffusivity for catalase, urease, aldolase, and alkaline phosphatase under the application of wide concentration ranges of substrates. Our single-molecule approach thus overcomes potential limitations and artifacts underscored by recent studies to show no enhanced diffusion in enzymatic reactions.

**Table of Contents artwork:** 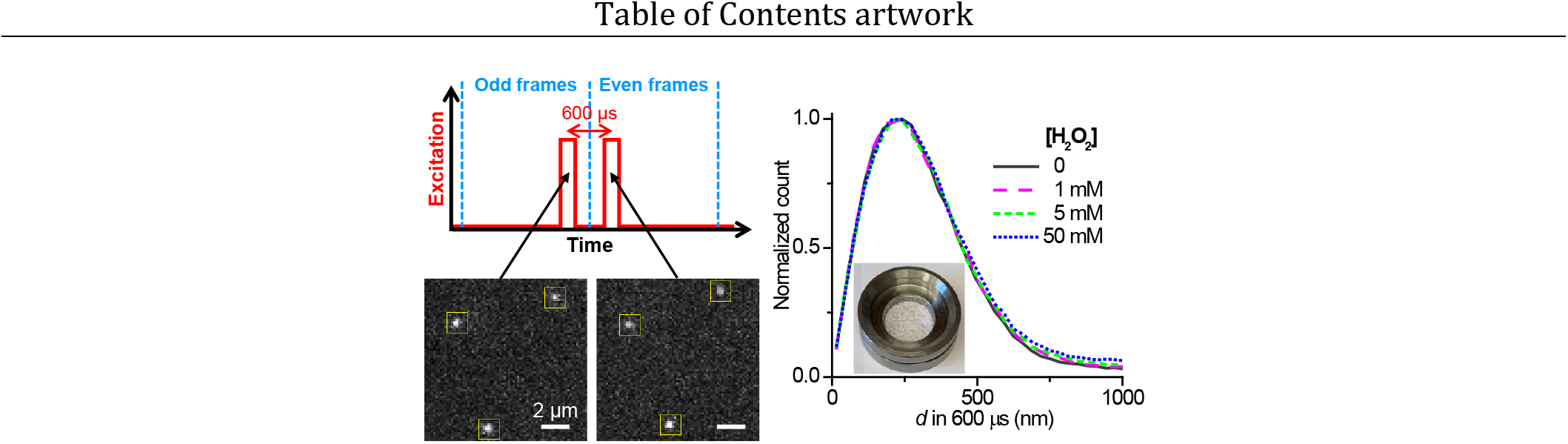

## TEXT

The recent reporting of accelerated diffusion of enzymes in catalytic reactions has generated keen research interest and heated debate.^1–8^ Measurements based on fluorescence correlation spectroscopy (FCS) have documented substantial (>~30%) increases in the diffusion coefficient *D* for multiple enzymes under catalytic turnovers.^1,2,9–14^ Several recent studies, however, emphasize FCS artifacts^3,15,16^ and report contradicting results. Dynamic light scattering (DLS) experiments, with low sensitivity for the very small sizes of proteins, also yields inconsistent results either supporting^13,14^ or refuting^17^ substrate-enhanced diffusion. Likewise, different nuclear magnetic resonance (NMR) experiments have separately reported the presence and absence of reaction-enhanced *D* for enzymes and small molecules,^18–21^ with unsettled debate on data interpretation.^8, 22,23^

Single-molecule experiments^24–26^ provide potential means to achieve precise *D* measurements by probing one molecule at a time. However, with expected *D*~30-50 μm^2^/s,^6^ the fast motions of typical enzymes make it difficult to detect freely diffusing single molecules. Using reducing-oxidizing chemicals that suppress fluorophore blinking, a recent study achieved long-time anti-Brownian electrokinetic (ABEL) single-molecule trapping of dye-labeled alkaline phosphatase (ALP) to show no change in *D* with the addition of its substrate.^16^ Meanwhile, single-molecule tracking has been reported for dye-labeled urease and ALP that were substantially slowed down by using viscous reagents as methylcellulose^27,28^ and glycerol^16,28^ or by tethering to supported lipid bilayers,^28^ which yielded contrasting results of no change versus up to 3-fold increases in *D* upon substrate addition. Thus, it remains difficult to quantify *D* through single-molecule measurements for molecules freely diffusing in regular buffers, whereas the addition of blinking-suppressing chemicals and viscous reagents complicates data interpretation.

We recently developed single-molecule displacement/diffusivity mapping (SM*d*M),^29,30^ a single-molecule imaging-based super-resolution method^31,32^ capable of determining molecular diffusivity. In SM*d*M, stroboscopic excitation pulses of *τ* <1 ms duration reduce motion-blur to capture snapshots of fast-moving molecules in the wide field. The excitation pulses are applied as pairs across tandem camera frames with a fixed Δ*t* <~1 ms interval (Figure 1a), so that images captured in the two frames (Figure 1b) can be compared to determine the transient displacements *d* of the observed molecules within the Δ*t* time window. Repeating this tandem excitation scheme then enables the accumulation of *d* over time for *D*-value determination.

**Figure 1.**
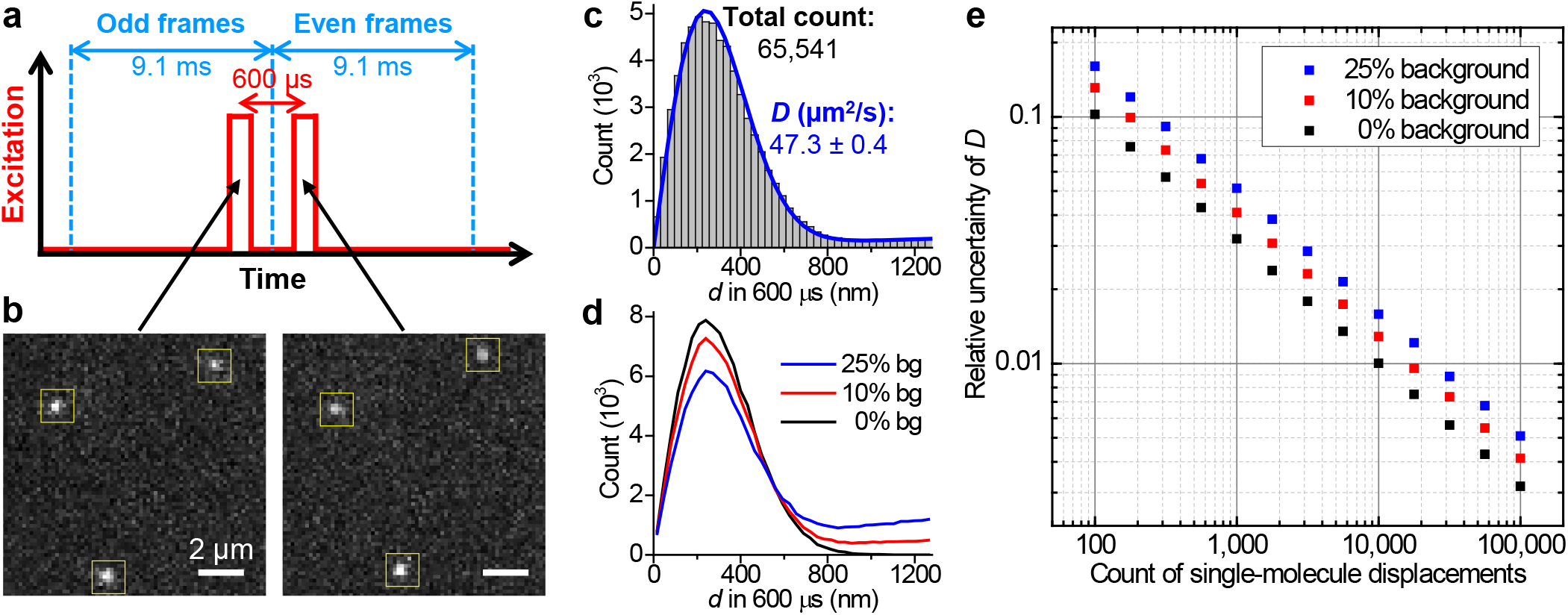
High-throughput statistics of unhindered single-molecule displacements. (a) Schematics: tandem excitation pulses of 300 μs duration are applied at Δ*t =* 600 μs center-to-center separations across paired frames of the recording EM-CCD camera. (b) Example single-molecule images captured in the paired frames, for Cy3B-labeled catalase freely diffusing in the PBS buffer, which enabled the determination of the transient displacements *d* of the 3 molecules marked by the yellow boxes in the Δ*t =* 600 μs time window. (c) Histogram: distribution of the 65,541 transient single-molecule displacements collected in 9 min by repeating the tandem excitation scheme 30,000 times. Blue line: MLE yielding a diffusion coefficient *D* of 47.3±0.4 μm^2^/s (95% confidence interval). (d) Example distributions of 100,000 simulated displacements, of which 0%, 10%, and 25% are backgrounds from molecules randomly entered the vicinity while the rest are 600-μs displacements with *D =* 50 μm^2^/s. (e) Relative uncertainty of *D* (*σ_D_/D*) from MLE results of the simulated data at different levels of backgrounds, as a function of the counts of displacements. The standard deviations *σD* were calculated from the differences between 5,000 rounds of simulations under each condition.

Whereas we previously focused on the high spatial resolution of SM*d*M to study intracellular diffusion heterogeneities,^29,30^ here we harness its capability for high single-molecule statistics to enable the precise quantification of *D* for enzymes freely diffusing in regular buffers. We thus formulate how precisions of better than ±1% may be achieved for *D* at the 95% confidence level, and then utilize this capability to show no changes in diffusivity for four highly contested enzymes under wide concentration ranges of substrates.

To optimize SM*d*M for the ~60% higher *D* versus the ~20-30 μm^2^/s values in our previous work on intracellular diffusion,^29^ we accordingly reduced the excitation pulse duration *τ* and pulse-to-pulse separation Δ*t* to 300 and 600 μs, respectively (Figure 1a and Figure S1). The use of Cy3B, a bright emitter,^33^ to label the enzymes enabled single-molecule detection with good signals (Figure 1b). By running the recording EM-CCD camera at 110 frames per second, ~10^4^ pairs of tandem frames were executed in minutes, from which >5×10^4^ single-molecule displacements were typically accumulated (Figure 1c). Maximum likelihood estimation (MLE) with a predefined model based on normal diffusion plus background (Methods) yielded good fits, from which *D* values were obtained with their 95% confidence intervals, *e.g.*, *D =* 47.3±0.4 μm^2^/s for Cy3B-labeled catalase in phosphate-buffered saline (PBS) (Figure 1c).

To define how the achievable precision in *D* depends on the count of single-molecule displacements *N*, we simulated 600-μs displacements *d* for molecules diffusing at *D =* 50 μm^2^/s. Different levels of backgrounds were further included, *i.e.*, assuming 0%, 10%, or 25% of the total counts were due to extraneous molecules that randomly entered the vicinity during Δ*t* (Figure 1d). MLE of the simulated data showed that with zero background, the relative uncertainty of *D* (*σ_D_/D*) was ~1/*√N*, so that 10% and 1% uncertainties were achieved with 100 and 10,000 counts of *d*, respectively (Figure 1e). This result is expected: each displacement provides an independent reporting of diffusivity, so the signal-to-noise ratio improves as ~*√N*. At background levels of 10% and 25%, 1% uncertainty in *D* was achieved with ~20,000 and ~30,000 counts of *d*, respectively. As our experimental data contained <10% background (compare Figure 1c and Figure 1d), ~0.5% relative uncertainty (hence an ~±1% bracket for the 95% confidence interval) was thus obtained (Figure 1e) with the typical >50,000 counts of *d* in our experiments, in agreement with our above MLE results on the experimental data (Figure 1c).

To verify whether the above high precisions would enable the experimental detection of small differences in *D*, we examined the diffusion of Cy3B-labeled catalase in PBS buffers containing varying amounts of glycerol. The addition of 5%, 10%, and 15% weight percentages of glycerol led to well-differentiated shifts in the measured *d* distribution (Figure 2a). The distributions remained well fitted with MLE (Figure 2b), which yielded ~10%, ~21%, and ~30% decreases in *D*, respectively (Figure 2c). Notably, two batches of samples measured a month apart yielded consistent *D* values, as well as good matches to trends predicted by the known glycerol concentration-dependent viscosity (Figure 2c). Among these results, the 2.5% glycerol sample showed ~5% reduction in *D* and was well differentiated from the glycerol-free sample (Figure 2c). Thus, our SMdM approach is well poised for detecting small differences in *D*.

**Figure 2.**
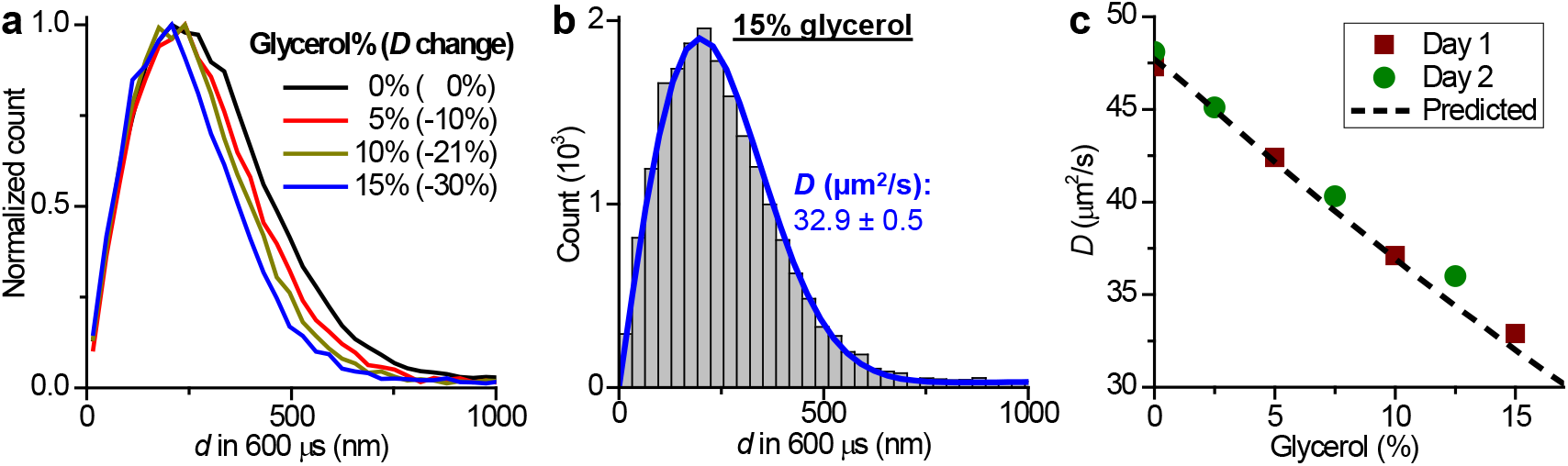
Precise determination of diffusivity *via* single-molecule statistics. (a) Normalized count distributions of the measured 600-μs single-molecule displacements for Cy3B-labeled catalase diffusing in PBS buffers containing 0%, 5%, 10%, and 15% glycerol. (b) Histogram: the count distribution with the 15% glycerol solution. Blue line: MLE yielding *D =* 32.9±0.5 μm^2^/s (95% confidence interval). (c) The measured *D* as a function of the glycerol percentage, for two batches of samples measured a month apart (squares and circles; 95% confidence intervals are comparable to the symbol sizes), compared to that is predicted based on the known viscosity of glycerol solutions (dash line).

For catalytic reactions, we started with catalase in the presence of its substrate H_2_O_2_, for which system FCS has reported ~30% increase in *D* with ~25 mM H_2_O_2_.^2,9^ SM*d*M statistics showed that the addition of 1, 5, and 50 mM H_2_O_2_ did not noticeably shift the single-molecule displacement distribution (Figure 3a). The occurrence of catalytic reaction was evident from the generation of oxygen bubbles, which were notable for samples with higher H_2_O_2_ concentrations (Figure 3a inset) and presumably led to the slightly increased backgrounds in the *d* distribution (tails in Figure 3a). Nonetheless, MLE yielded good fits (Figure 3b), and the resultant *D* values showed minimal variations in the range of 46.5-48.0 μm^2^/s between the different substrate concentrations (Figure 3c). The ~±1.5% variations, while slightly larger than our ~±1% bracket for 95% confidence, which may be attributed to uncontrolled experimental factors, are significantly smaller than the ~30% changes reported in previous FCS studies.^2,9^

**Figure 3.**
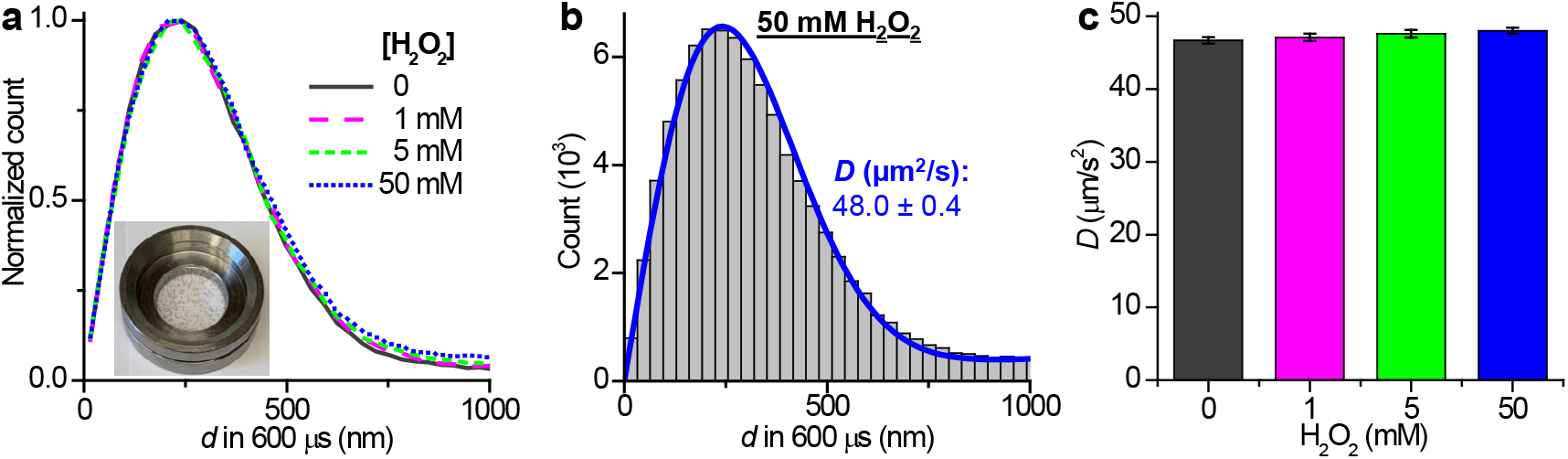
No substantial change in single-molecule displacements observed for catalase under turnover. (a) Normalized count distributions of the measured 600-μs single-molecule displacements for Cy3B-labeled catalase diffusing in PBS buffers with H_2_O_2_ added at 0, 1, 5, and 50 mM. Inset: photo of a sample with 50 mM H_2_O_2_, showing the notable generation of bubbles from the catalytic reaction. (b) Histogram: the count distribution with the 50 mM H_2_O_2_ solution. Blue line: MLE yielding *D =* 48.0±0.4 μm^2^/s (95% confidence interval). (c) MLE-determined *D* at the different substrate concentrations. Error bars: 95% confidence intervals.

We next examined aldolase, for which system previous FCS experiments have either reported ~30% increase^10^ or no change^14^ of *D* in the presence of ~1 mM of its substrate, fructose 1,6-bisphosphate (FBP), whereas DLS^13,17^ and NMR^19^ experiments have indicated no *D* enhancement over broad substrate ranges. SM*d*M showed virtually identical distributions for the 600-μs single-molecule displacements with and without the addition of 0.1-10 mM of FBP (Figure 4a). MLE hence yielded comparable *D* values of 44.7-46.8 μm^2^/s for the six conditions with no noticeable trend on the substrate concentration (Figure 4d).

**Figure 4.**
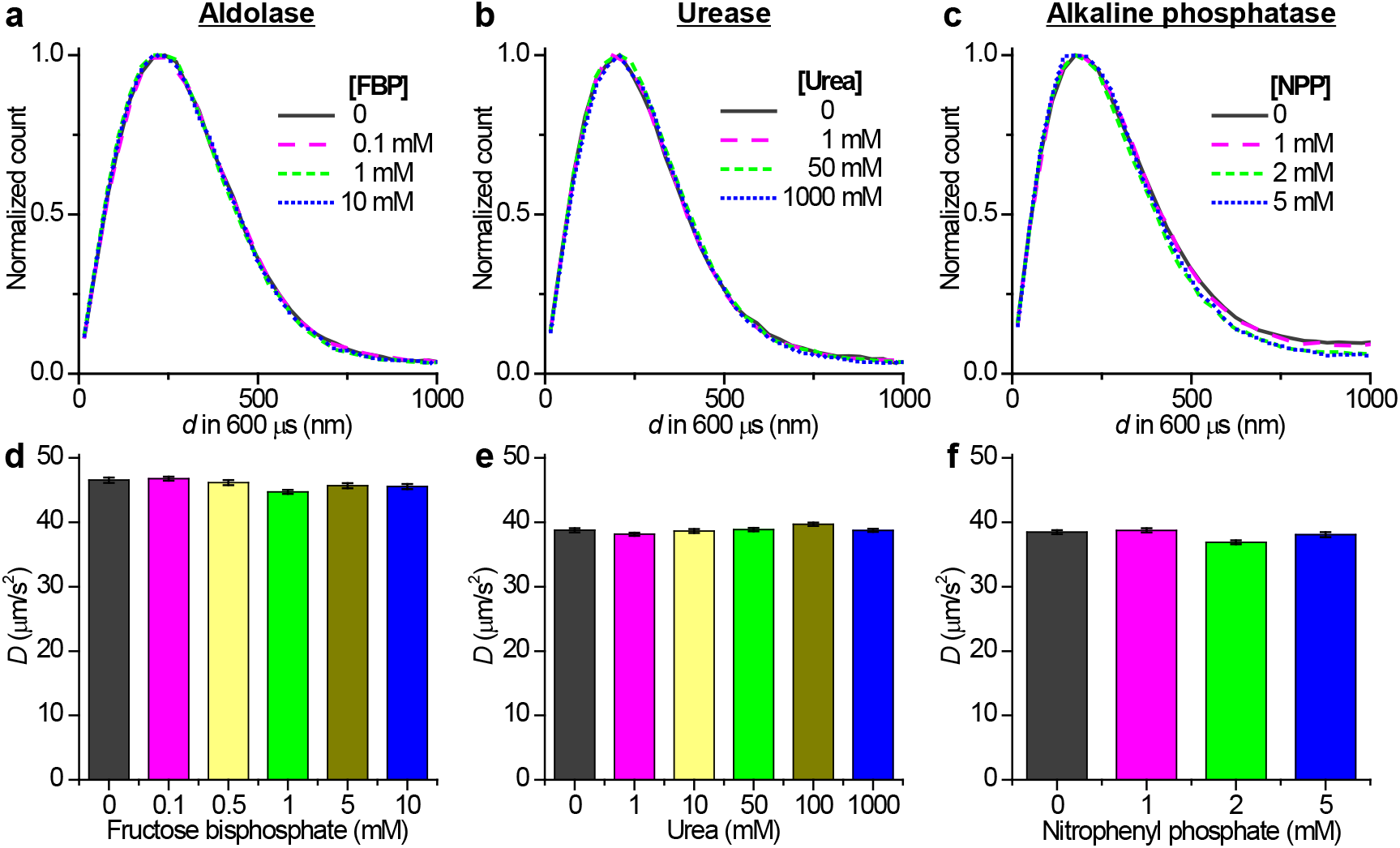
Results on aldolase, urease, and alkaline phosphatase also show no changes in single-molecule displacements under catalytic turnover. (a-c) Normalized distributions of the measured 600-μs single-molecule displacements for Cy3B-labeled aldolase (a), urease (b), and alkaline phosphatase (c), in PBS (a,b) and Tris (c) buffers with and without the addition of different concentrations of the corresponding substrates: fructose 1,6-bisphosphate (a), urea (b), and *p*-nitrophenyl phosphate (c). (d-f) MLE-determined *D* values for the samples with different substrate concentrations. Error bars: 95% confidence intervals.

Similarly, we detected effectively invariant single-molecule displacement distributions for urease when its substrate, urea, was added in the wide range of 1-1000 mM (Figure 4b). The resultant consistent *D* values of 38.2-39.7 μm^2^/s across six substrate concentrations (Figure 4e) thus contrasted with previous reports of up to ~40% enhancement in *D* with FCS,^1, 2,11,13,14^ as well as recent results reporting ~3-fold *D* enhancement based on the tracking of single urease molecules that diffuse orders of magnitudes slower in highly crowded or membrane-tethered systems.^27,28^

We next turn to ALP, for which FCS has initially reported up to 80% increase in *D* with the addition of 2.6 mM of its substrate *p*-nitrophenyl phosphate (NPP).^2^ However, recent FCS experiments,^3, 14,16^ as well as single-molecule trapping in blinking-suppressed buffers and tracking in glycerol-slowed solutions^16^ indicate no reaction-induced *D* enhancement. Using a 5 mM pH = 8.0 Tris-HCl buffer containing 2 mM MgCl_2_ and 50 μM ZnCl_2_, the preferred working condition for ALP,^2^ SM*d*M showed that the distribution of single-molecule displacements and hence *D* values did not alter substantially with the addition of 1-5 mM NPP (Figure 4cf).

Together, with the high precision achieved through the high single-molecule statistics of SM*d*M, our results consistently showed unvarying *D* for four enzymes that have been reported as exhibiting enhanced diffusion under catalytic turnovers. These results, together with recent experiments that reexamine artifacts due to photophysical processes as fluorescence quenching and blinking,^3,15,16^ underscore the difficulties with quantifying *D* through FCS. Although blinking-suppressing buffers partly alleviate these issues while also enabling single-molecule trapping for *D* measurements,^16^ the addition of reducing and oxidizing chemicals complicates data interpretation. Meanwhile, the high *D* values of enzymes have largely limited single-molecule tracking to systems in which viscous reagents as methylcellulose^27,28^ and glycerol^16,28^ are added to slow down molecular motions.

In contrast, with SM*d*M, here we were able to work with the diffusion of enzymes in regular buffers without the addition of extraneous components that may potentially affect the system: As SM*d*M captured and accumulated the transient (600 μs) displacements of single molecules, it removed the needs to establish long-term fluorophore photostability or to impede diffusion to permit multi-frame tracking. Whereas in this study we have employed MLE with a model based on normal diffusion to extract *D*, we emphasize that our results are not tied to any particular model. As shown from our control experiments, small changes in diffusivity would lead to notable shifts in the distribution of single-molecule displacements: our observation of highly invariant distributions under different substrate concentrations indicated no changes in diffusivity, independent of the underlying model. Our results thus support recent analysis that reaction-facilitated diffusion vanishes at the molecular scale.^8^ The application of the tools developed in this work to other systems in which precise *D* values are important, potentially including the even faster diffusion of smaller molecules, awaits future efforts.

## ACKNOWLEDGMENTS

This work was supported by the National Science Foundation (CHE-1554717), the National Institute of General Medical Sciences of the National Institutes of Health (DP2GM132681), the Beckman Young Investigator Program, and the Packard Fellowships for Science and Engineering, to K.X. K.X. is a Chan Zuckerberg Biohub investigator.

## Supporting Information

### Materials and Methods

#### Fluorescent labeling of enzymes

Aldolase from rabbit muscle, alkaline phosphatase (ALP) from bovine intestinal mucosal, catalase from bovine liver, and urease (type III) from jack bean were purchased from Sigma-Aldrich (A8811, P7640, C30, U1500). The enzymes were labeled with Cy3B NHS (*N*-hydroxysuccimidyl) ester (Cytiva, PA63101) in 0.1 M NaHCO3 at a ~10:1 dye-to-protein ratio and reacted for 4 h at room temperature. Excess dye was removed using Amicon 3k MWCO Centrifugal Filters (Millipore, UFC500396). Absorbances at 280 and 560 nm, as measured by a NanoDrop 2000c spectrometer (ThermoFisher), showed that the labeled product had 0.5-2 dyes per protein with a protein concentration of 0.1-1.0 mg/mL. The labeled aldolase, catalase, and urease were kept in Dulbecco’s Phosphate-Buffered Saline (DPBS) on ice. The labeled ALP was kept in 5 mM Tris pH = 8.0, 2 mM MgCl_2_, 50 μM ZnCl_2_ on ice. Single-molecule experiments were performed on the same day of labeling.

#### Sample preparation

25-mm diameter, #1.5 coverslips were treated in a heated 3:1 H_2_SO_4_ (98%) and H_2_O_2_ (30%) mixture for 45 minutes, rinsed until neutral using Milli-Q water, and then dried using N_2_ gas. Coverslips were functionalized with 2 mM methoxy PEG silane (5 kDa, PG1-SL-5k, Nanocs) in 95% ethanol/water for 30 minutes in an ethanol vapor. Coverslips were rinsed and sonicated for 5 minutes in Milli-Q water before being mounted into an imaging chamber (ThermoFisher, A7816). Cy3B-labeled enzymes were diluted to ~200 pM into the buffer (DPBS for urease, catalase, and aldolase; 5 mM Tris pH = 8.0, 2 mM MgCl_2_, 50 μM ZnCl_2_ for ALP) with or without the addition of substrates at different concentrations, and then immediately imaged. Substrates used: D-fructose 1,6-bisphosphate trisodium salt hydrate (Sigma-Aldrich, F6803), *p*-nitrophenyl phosphate (Sigma-Aldrich; 4876), hydrogen peroxide 30% (VWR, BDH7742-1), and urea (EMD, UX0065-1).

#### SM*d*M of enzymes

SM*d*M was performed on a Nikon Ti-E inverted fluorescent microscope, as described previously,^1^ but with substantially reduced excitation pulse duration *τ* and pulse-to-pulse separation Δ*t*. Briefly, a 561 nm laser (OBIS 561 LS, Coherent, 165 mW) was focused to the back focal plane of an oil-immersion objective lens (CFI Plan Apochromat Lamda 100X Oil, NA = 1.45) to enter the sample slightly below the critical angle of the glass-solution interface, so that it illuminated a few micrometers into the solution. The focal plane was set at ~2 μm away from the glass surface into the solution, which was maintained by the native Nikon Perfect Focus System (PFS) on the microscope during the experiment. A multifunction I/O board (PCI-6733, National Instruments) detected the exposure timing signal of the recording EM-CCD camera (iXon Ultra 897, Andor), and accordingly modulated the emitting power of the 561 nm excitation laser. Paired pulses of τ = 300 μs duration and Δ*t =* 600 μs center-to-center separation were thus repeatedly applied across tandem camera frames, with the timing waveforms confirmed by a GW Instek GDS-1054B oscilloscope (Figure S1). Typical runs executed 1.7-3.0×10^4^ pairs of tandem frames at 110 frames per second (fps), thus data-collection times of 5-9 min. Fluorescence emission was filtered by a longpass (ET575lp, Chroma) and a bandpass filter (ET605/70m, Chroma).

#### Data analysis

SM*d*M data were analyzed as described previously.^1^ Briefly, single molecules were first super-localized in each recorded frame (*e.g.*, Figure 1b), and then their two-dimensional displacements *d* between the paired frames were calculated with a search radius *R* of 1.3 μm. The accumulated >5×10^4^ single-molecule displacements in each run were pooled together for plotting of distribution (*e.g.*, Figure 1c) and fitting to the below model:^1^

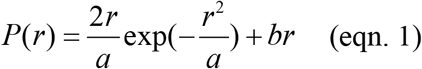

Here *a =* 4*D*Δ*t*, and *b* accounts for a uniform background due to extraneous molecules that randomly diffuse into the search radius during Δ*t*.^1^ MLE with MATLAB returned 95% confidence intervals for *a*, which were converted to *D* results with our fixed Δ*t =* 600 μs in the experiment.

#### Simulation

Monte Carlo simulation was performed with MATLAB, borrowing components from our recent simulation work on more complicated systems.^2^ Single-molecule displacements in Δ*t =* 600 μs were simulated for different total counts based on normal diffusion (two-dimensional random walk)^3^ with *D =* 50 μm/s, with the addition of Gaussian blurs of *σ =* 20 nm for both the starting and ending positions to further simulate the uncertainties in localizing the single molecules. Moreover, we allowed extraneous molecules to randomly enter the simulated area with a uniform spatial density to contaminate 10% or 25% of the total counts. The resultant simulated single-molecule displacement data (see example distributions in Figure 1d) were fed as input for the MLE determination of *D* based on eqn. 1 as described above. The simulation under each condition was repeated 5,000 times, so that the differences in the extracted *D* values between the 5,000 runs were used to calculate the standard deviation *σD*. The relative uncertainty *σD*/*D* was then plotted as a function of the count of simulated displacements (Figure 1e).

## Supplementary Figures

**Figure S1.**
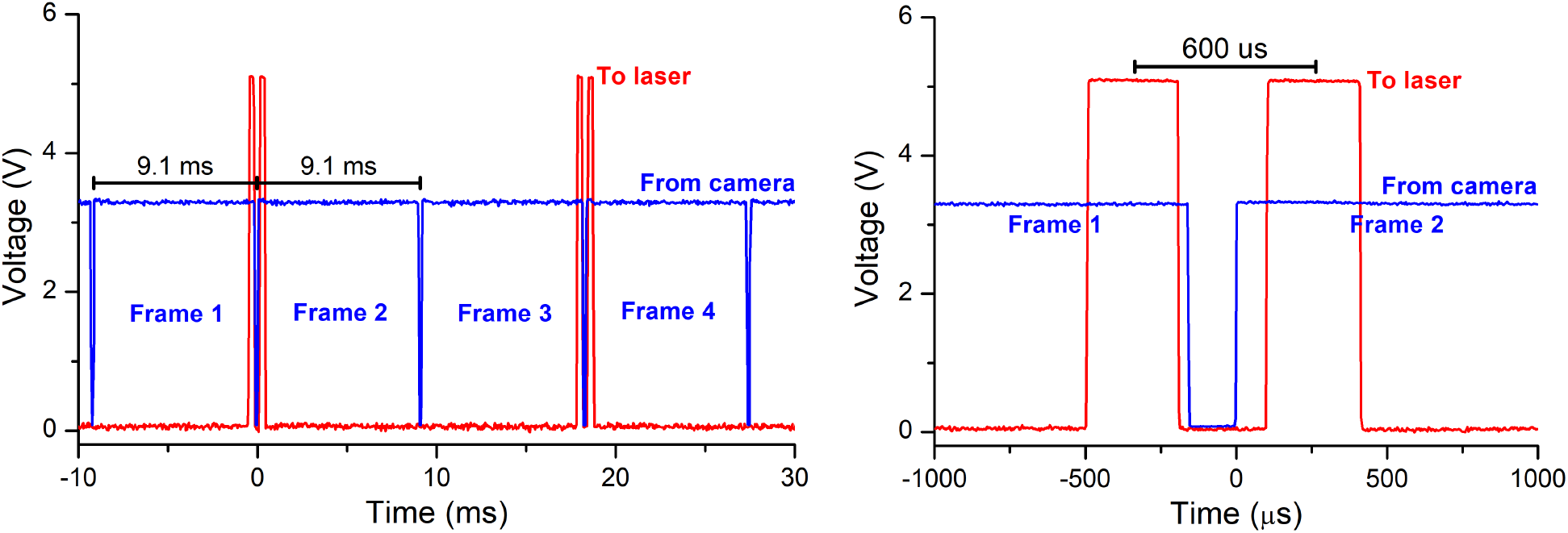
Timing waveforms used in this work. Left panel: zoom-out for four consecutive camera frames. Right panel: zoom-in for the tandem pulses across two frames. Blue traces: the “fire” TTL signal from the EM-CCD camera (Andor iXon Ultra 897), which ran at 110 fps (9.1 ms/frame). 3.3 V corresponds to when exposure occurred, whereas 0 V corresponds to the dead time between frames. Red traces: output from the PCI-6733 card, which drives the 561 nm excitation laser between the off (0 V) and on (5 V) states *via* digital modulation. A pair of pulses each of *τ =* 300 μs duration are applied across the two frames at a Δ*t =* 600 μs center-to-center separation, avoiding the dead time of the camera.

